# Consensus-seeking and conflict-resolving ---an fMRI study on college couples’ shopping interaction

**DOI:** 10.1101/2020.03.11.986786

**Authors:** HanShin Jo, Chiu-Yueh Chen, Der-Yow Chen, Ming-Hung Weng, Chun-Chia Kung

## Abstract

One of the typical campus scenes is the social interaction between college couples, and the lesson couples must keep learning is to adapt to each other. This fMRI study investigated the shopping interactions of 30 college couples, one lying inside and the other outside the scanner, beholding the same item from two connected PCs, making preference ratings and subsequent buy/not-buy decisions. The behavioral results showed the clear modulation of significant others’ preferences onto one’s own decisions, and the contrast of the “shop-together vs. shop-alone”, and the “congruent (both liked or disliked the item, 68%) vs. incongruent (one liked but the other disliked, and vice versa)” together trials, both revealed bilateral temporal parietal junction (TPJ) among other reward-related regions, likely reflecting mentalizing during preference harmony. Moreover, when contrasting “own-high/other-low vs. own-low/other-high” incongruent trials, left anterior inferior parietal lobule (l-aIPL) was parametrically mapped, and the “yield (e.g., own-high/not-buy) vs. insist (e.g., own-low/not-buy)” modulation further revealed left lateral-IPL (l-lIPL), together with left TPJ forming a local social decision network that was further constrained by the mediation analysis among left TPJ-lIPL-aIPL. In sum, these results exemplify, via the two-person fMRI, the neural substrate of shopping interactions between couples.

## Introduction

A matured society relies on the good functioning of her building blocks--families, in various domains and in different stages. For example, newlyweds have to gather savings or loans for their future home; first time parents prefer pricier, though not always healthier, option^1^ when purchasing children-related items^2^, and not to mention the later education funds. All these are contingent on the readiness of young adults sensitive to the needs of, and the willingness to sacrifice one’s own interest for the benefit of, significant others. As a precursor, romantic young couples have not just been shown to prefer making important decisions jointly than individually^3–5^, they also prioritize the interest of significant others than that of oneself^6^.

When making group economic decisions, one has to put him/herself in another’s perspective, a process commonly called theory-of-mind, or TOM, in simulating other-oriented decisions^7,8^. On one hand, behavioral research has demonstrated that consumer’s purchase decisions are highly influenced by others (e.g., feedbacks from peers or online customers)^9–11^ and social/cognitive/emotional factors^12–15^; on the other, neuroscientists also uncovered the neural substrates of TOM in various forms of social cognition^16,17^: for example, one earlier meta-analysis study^18^ listed the neural substrates of TOM, which include bilateral temporal parietal junction (TPJ), posterior superior temporal sulcus (pSTS), and medial prefrontal cortex (mPFC), as the core unit of TOM. In addition, precuneus, temporal lobe, and inferior frontal gyri are selectively associated with task-specific TOM contexts^18^. More importantly, the brain regions surrounding TPJ, including the inferior parietal lobule (IPL) and superior temporal sulcus (STS/STG), are also heavily involved in various contexts of social inferences^19,20^, such as empathy concerns^21,22^, prosocial behaviors^23,24^, cooperations^25^, social compliances and conformities^26–28^. Although the above-mentioned brain regions are concorted in various paradigms and affective/cognitive contexts, their specific roles and functional interactions in everyday cognition remain less discussed.

To address these concerns, the present event-related fMRI study explores the neural substrates, as well as their functional interactions, of romantic college couples in an interactive shopping task (Figure. 1). During the experiment, participants gave separate preference ratings on each presented product, and their preferences were revealed to each other ⅔ of the time, to allow modulation of significant other’s evaluation before one’s own shopping decisions. This paradigm created various scenarios like: ‘shop together’ versus ‘shop alone’, which we assume would contrast TOM-related regions (such as bilateral TPJ^29^), and ‘preference (in)congruence’ (the differences between couples’ item ratings), and more interestingly, in cases of preference incongruent scenarios: subsequent decision changes (i.e., yielding to significant other’s evaluation) or not (i.e., insisting on one’s own preference). Building on the extant neuroimaging literature of empathic choices or conformity under moral concerns^30,31^ or social norms^24,32–34^, the mPFC, TPJ, dorsal anterior cingulate cortex (dACC), nucleus accumbens (NAcc), and insular cortex are all relevant brain areas expected to be activated under various parametric settings. Although to our knowledge, the present study is the first to implement real-life interactions between college couples’ shopping behaviors with fMRI, we still expect the findings, primarily via general linear model, parametric modulation, functional connectivity (e.g., psychophysiological interactions, aka., PPI^35^) and confirmatory effective connectivity (subject-level mediation analysis)^36^, to be largely comparable to the earlier results through self/other evaluations^26,37,38^, conflict resolving^39,40^, and/or consensus reaching^41–43^.

**Figure 1.**
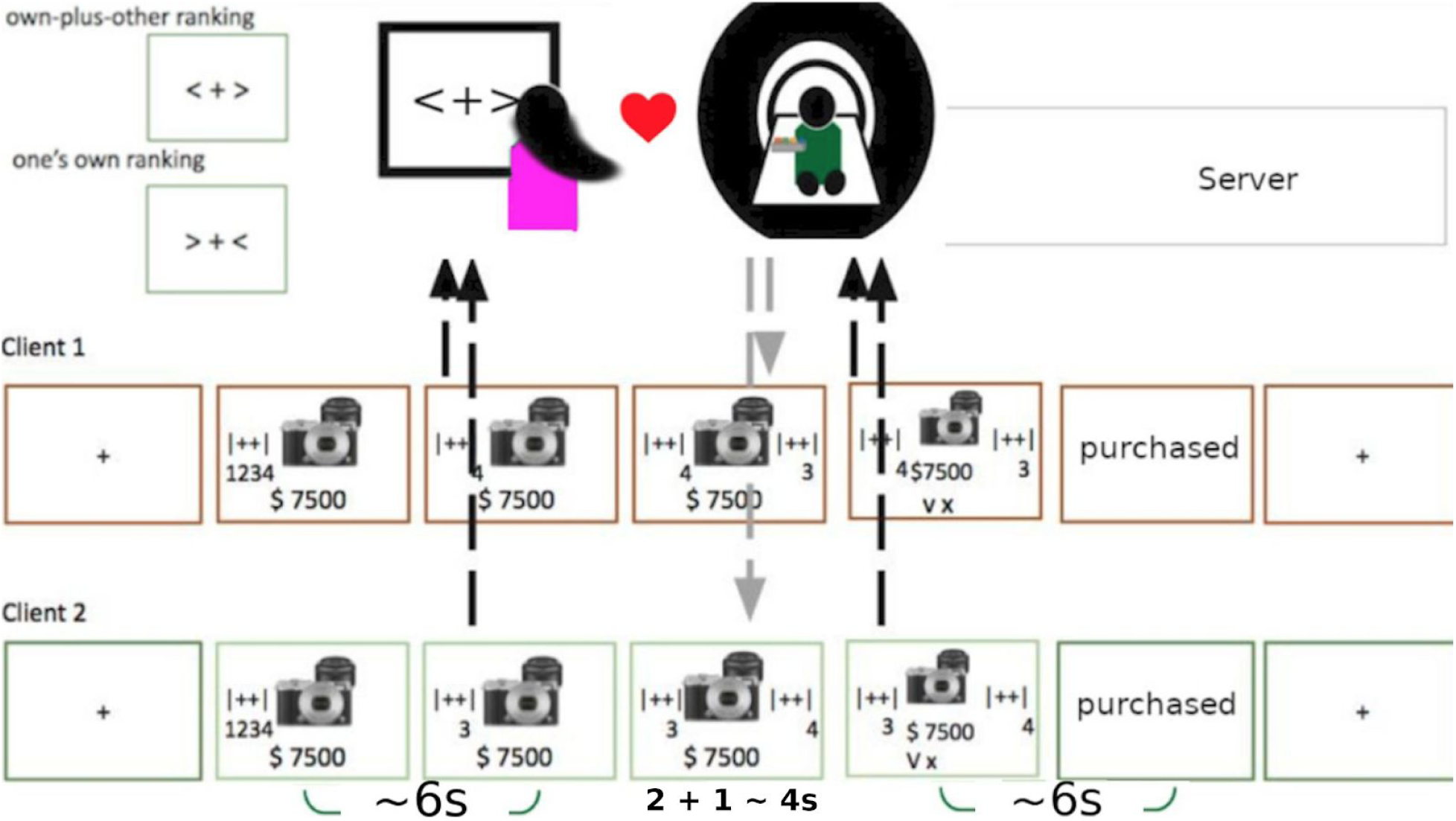
Experimental design. Two types of shopping contexts were designed to examine how significant others’ preferences affect one’s own shopping decisions. After 1 sec of fixation cross with arrows indicating alone/together shopping at the beginning of each trial, stimulus was presented with the price. During 6 secs of evaluation time, participants give preference ratings from 1 (dislike most) to 4 (like most). Then the two screens would show either one’s own rating or own-plus-other ratings for a jittered interval (3 ∼ 6 secs). Subsequently, both a ‘checked’ and a ‘X’ mark would appear in the lowest part of the screen, cueing the beginning for response period (up to 6s). Each participant’s responses would be recorded and displayed independently by a ‘purchased’ or ‘not-purchased’ sign.

## Results

### Behavioral results

Out of the 2102 trials collapsing across 30 participants, 68% of the preference ratings were congruent across couples: either both high (pressing 3 or 4) or both low (1 or 2), and the remaining incongruent trials (32%) were one high (3 or 4) and the other’s low (1 or 2), and vice versa. In addition, in 62.5% of incongruent evaluations (or 20%, = 62.5%*32% of all trials), participants’ shopping decisions were consistent with their *own* preferences (or the so-called *‘insisting’*, meaning the choice was done *despite of* significant other’s opposite opinion), and the remaining 37.5% incongruent trials (or 12% of all trials) participants were ‘*yielding*’ (meaning to change one’s own decision against one’s own preference) to significant other’s preferences.

Figure 2 shows the behavioral results of shopping ratios under ‘shop alone’ (left) vs. ‘shop together’ (right) conditions. In shop-alone conditions, one’s own buy ratios were positively correlated (r=0.65, p < 0.01) with one’s own preference ratings. This is not surprising, since the low preference (1 & 2) reflects item’s low likability, and therefore the low buying rates (and vice versa). In contrast, when comparing Fig. 2 left with 2 right, other than the similar correlation between one’s own preference and buying ratio, one can additionally observe the clear modulation of significant other’s preferences: when compared to the ‘shop-alone’ baseline (Fig. 2 left), not only were significant other’s ratings both decreased/increased one’s buying ratio when both gave low/high ratings, but also the similar modulation happened when both people gave opposite preference ratings: people yielded more in times of conflicts by increasing the buying ratio when facing significant other’s higher rating, and decreasing when facing the other’s lower rating. Out of the overall shopping ratio across all preferences (36%, SD = 0.406), the two-way ANOVA showed that buying ratio was significantly different between shop-alone vs. shop-together (across other’s preference: 1 (F(4,171) = 3.85, p < 0.005); 2 (F(4,184) = 7.83, p < 10^−4^); 3 (F(4,184) = 16.77, p < 10^−5^); 4 (F(4,145) = 6.01, p < 10^−4^). see Figure 2). The female and male participants did not differ significantly in preference nor in shopping decisions (both p > .05), reflecting a possible social desirability effect.

**Figure 2.**
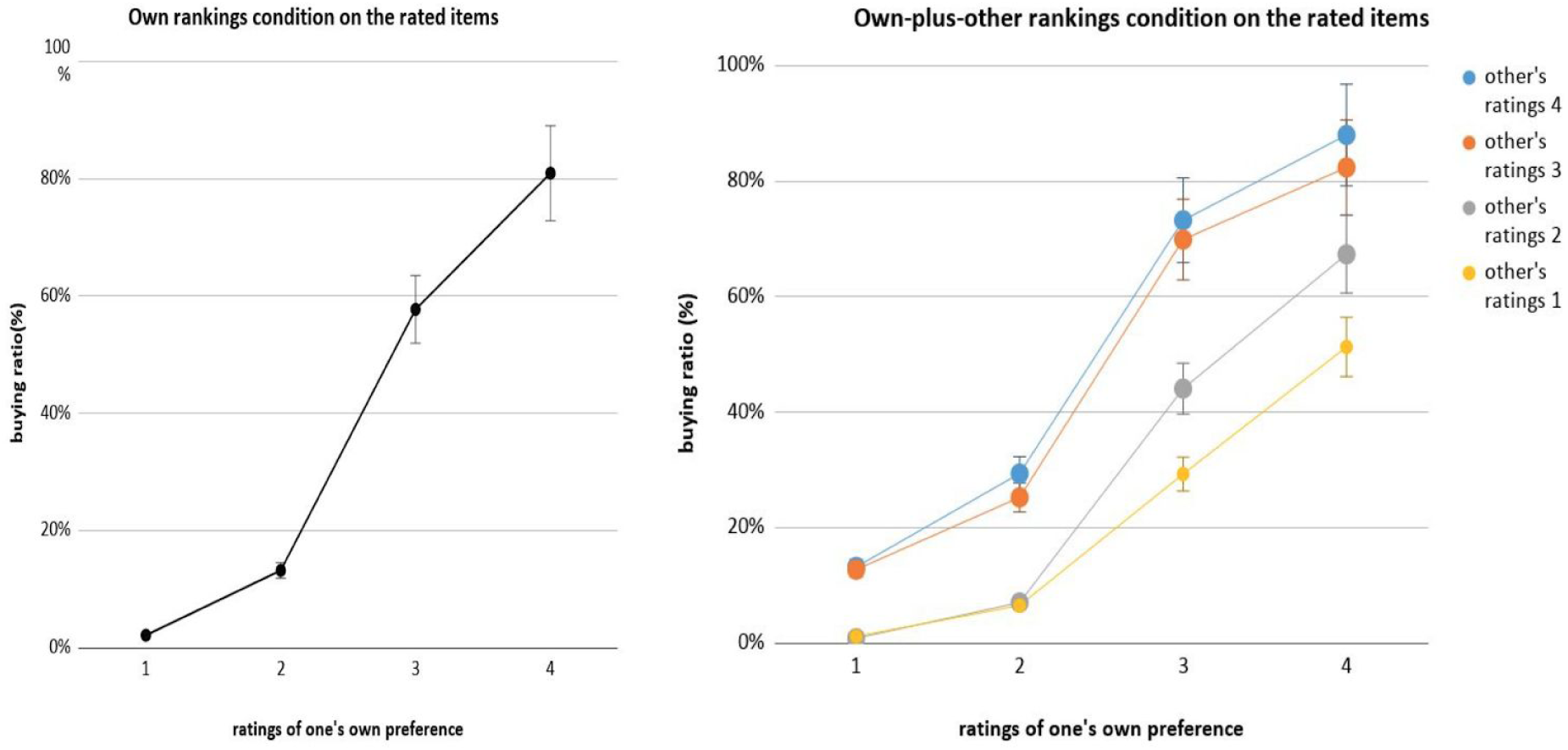
Behavioral analysis results. The comparison of buying ratio between alone and together condition, the modulatory effect of significant other’s preference was observed by increasing one’s buying decision significantly when partner’s preference was high (3 or 4), and decreasing one’s buying decision when partner’s preference was low (1 or 2) compared to shopping alone. The two-way ANOVA showed significant differences between shop-alone vs. shop-together as follows: across other’s preference: 1 (F(4,171) = 3.85, p < 0.005); 2 (F(4,184) = 7.83, p < 10^−4^); 3 (F(4,184) = 16.77, p < 10^−5^); 4 (F(4,145) = 6.01, p < 10^−4^. The female and maleparticipants did not differ significantly in preference nor in shopping decisions (both p > .05).

The logistic regression revealed that both own and other’s preference ratings were significant predictors for buy/not-buy decisions (alone-own: b = 2.48, p < 10^−3^; together-own: b = 2.80, p < 10^−4^; together-other: b = 1.74, p < 10^−3^), as well as the preference difference and interaction between the two (own-other rank differences: b = -0.97, p < 10^−3^; interaction between own and other rank: b = -0.29, p < 0.01). The negative significance meant that the modulation of significant other’s preference was decreasing as the self preference increased. Additionally, price and age were identified as significant predictors in both alone and together conditions, suggesting that pricier items and older (maturier) participants were both shopped and shopping less (See Supplementary Table S1).

## fMRI Results

### General linear model of interactive shopping task

In GLM analysis, the brain regions revealed in the “together vs. alone trials” contrast were: bilateral temporal parietal junction (TPJ), superior temporal gyrus (STG), ventral striatum, posterior cingulate cortex (PCC), and dorso-medial prefrontal cortex (dmPFC), all higher activated in the former (together) condition (see Figure 3 left). The “incongruent vs. congruent” contrast, in turn, was defined by “both high or both low” vs. “either A-high-B-low or A-low-B-high” (low being 1/2 and high being 3/4). This contrast revealed bilateral TPJ, posterior insula, putamen, and PCC/precuneus as more engaged in congruent (than incongruent), meanwhile, anterior insula (AI) and superior frontal gyrus (blue region) more in incongruent trials (see Figure 3 right). Since the majority (68%) of the together trials were congruent type, finding bilateral TPJ in both contrasts is not surprising. Whole brain cluster table has been provided as supplementary information (see Supplementary Table S2).

**Figure 3.**
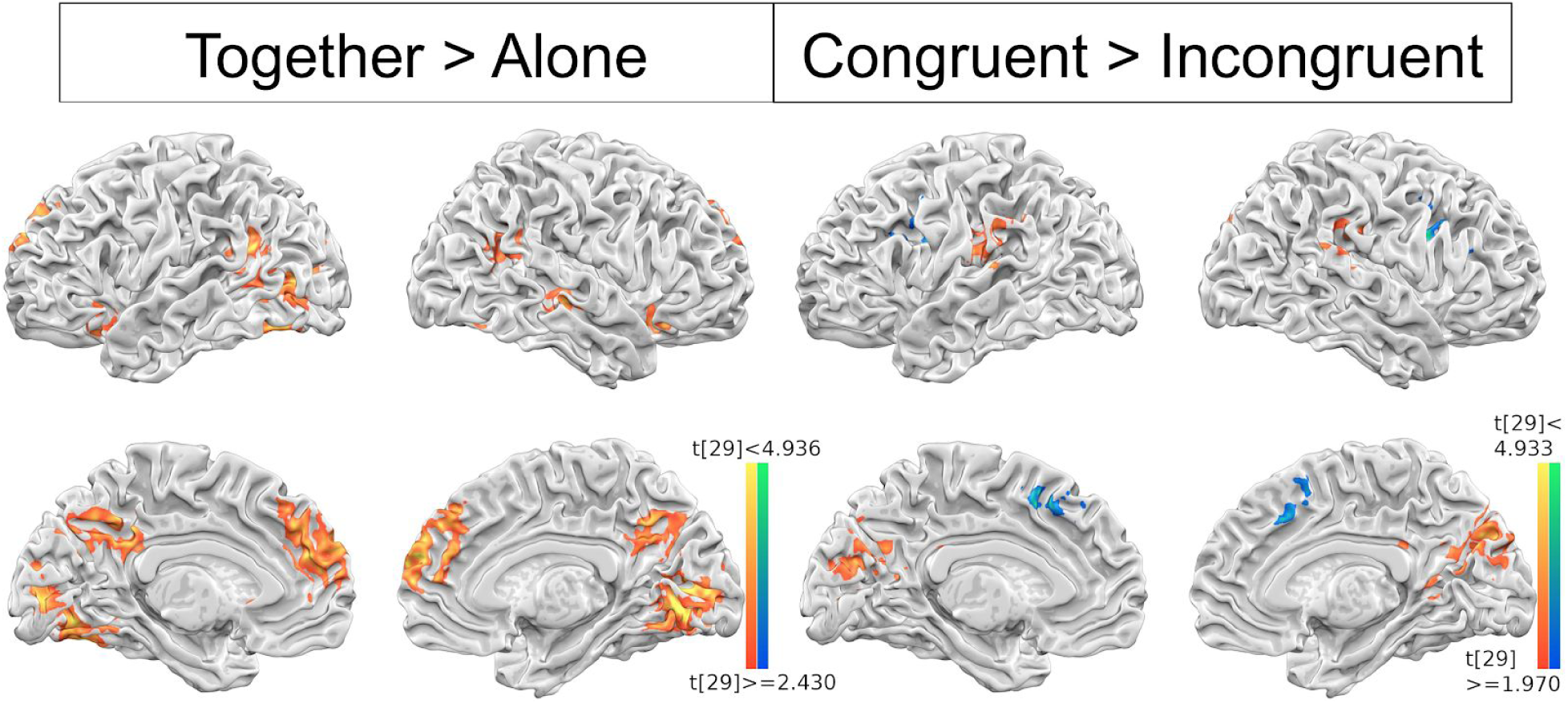
GLM analysis results. The brain regions engaged in together > alone (left) and congruent > incongruent (right) contrasts. The bilateral TPJ, STG, ventral striatum, PCC, and dmPFC were commonly involved in shopping together than alone, in turn, ‘congruent > incongruent’ contrast defined by chosen preferences in two categories (high = 3,4/low = 1,2) revealed brain regions such as bilateral TPJ, insular, putamen, and PCC/precuneus for congruent (same rating categories), and SFG for incongruent (different rating categories) condition. The contrasts are thresholded at p < 0.01 (Left) and p < 0.05 (Right), and FWE corrected at p < 0.05.

### Decision making under incongruent situation: High vs. Low preference and Insisting vs. Yielding

The next analysis was more focused on resolving incongruent situations. First, the incongruent situation could be further divided into when own preference is high, but partner’s preference is low and vice versa. To address two different underlying psychological responses, we performed additional parametric GLM with own high preference (/other low preference) to low preference (/other high preference) as a linear modulatory regressor. The left anterior IPL was identified as a brain regions playing modulatory role for self-high/other-low, whereas the PCC activation represented increasing conflict of self-low/other-high case (figure 4 left).

**Figure 4.**
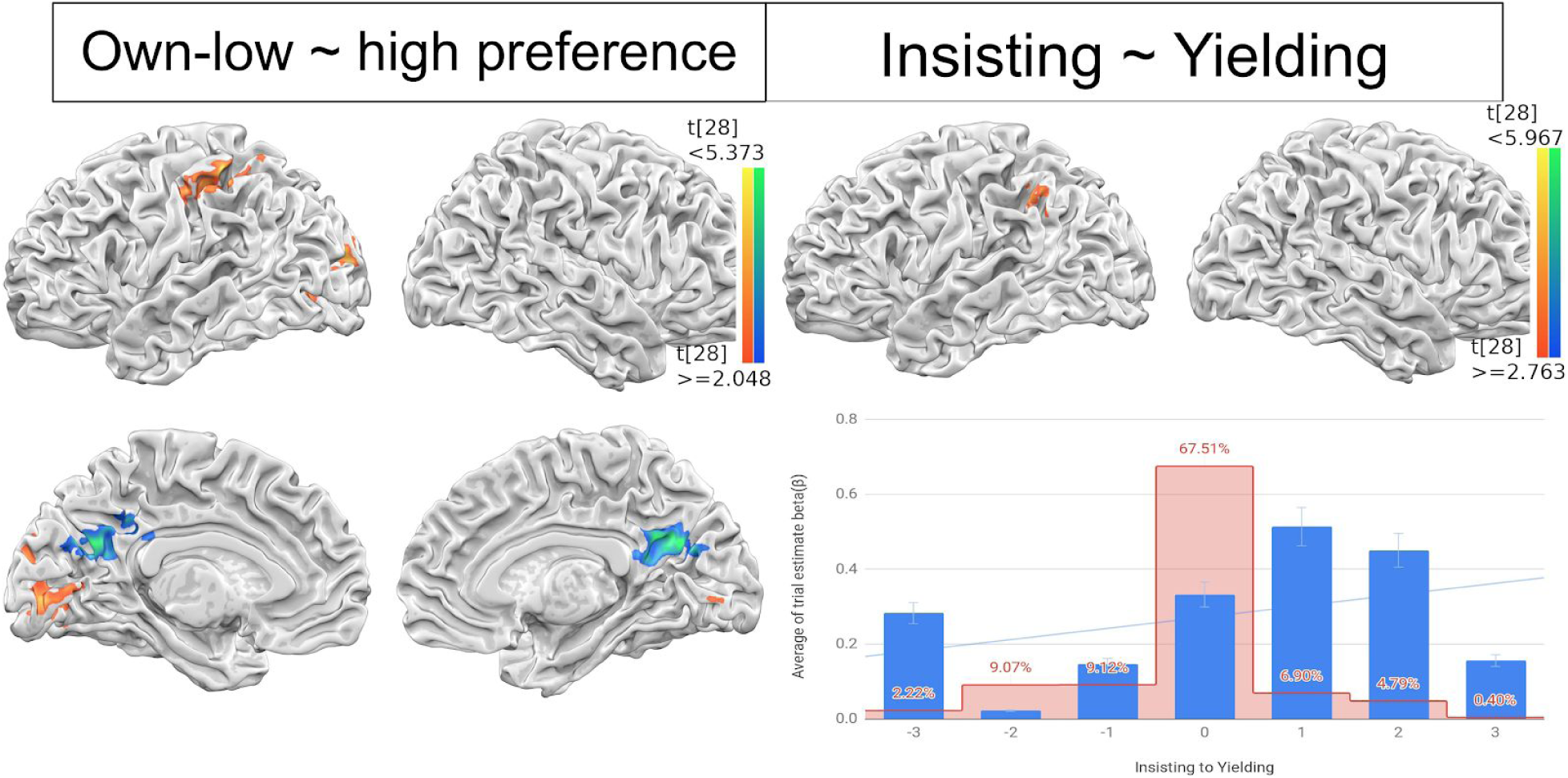
Parametric modulation results. The parametric modulation of preference differences (left) and decision changes (right) in incongruent condition. The preference differences were parameterized by subtracting preferences between own and others in own-high/other-low, recruiting left anterior IPL, or own-low/other-high cases, in PCC. The decision changes were further encoded by multiplying the decision (buy/not-buy) toward own preference (insisting) or other preference (yielding) to preference differences, only the left lateral IPL activations were positively correlated. The trial estimate beta coefficient (*β)* extracted from the left IPL were averaged and plotted across insisting (−3) to yielding (3) to show parametric relations (bottom right). The surface map of preference differences was thresholded at p < 0.05, FWE corrected and decision change was thresholded at p < 0.01, k = 20 (uncorrected).

The preference scores were further encoded as the decision change value, representing the partner checked the other’s preference and then decided to change their mind to reach agreement (yielding/giving in) or insist their mind with own preference (insisting). The decision changes as a parametric regressor, parametric GLM was computed. The results revealed that left IPL has positively correlated with increasing decision changes toward others preference (figure 4 right top). The linear regression was drawn to describe the relationship between left IPL and decision changes. The left IPL voxel trial estimated beta was extracted and average beta coefficient was shown positive relationship with corresponding decision change value (figure 4 right bottom, due to the small number of trials where 3 steps of behavioral change take place, without the 3 steps parameters, the regression was more significant).

### The functional and effective connectivity of congruency-driven decision processing

In the functional connectivity, or Psycho-Physiological Interaction (PPI), analysis, the 4 different seeds: bilateral TPJ, and left anterior, and left lateral IPL, were chosen to reveal their corresponding brain regions differentiating between incongruent and congruent item-evaluation contexts. The PPI results showed that the left TPJ and left lateral IPL seeds are both functionally connected more (in incongruent than congruent condition) to bilateral mPFC, dACC, caudate, insular, and inferior frontal gyrus (IFG); whereas dmPFC and middle frontal gyrus are specifically more connected to l-TPJ seed, and left mPFC and superior frontal gyrus more to l-lIPL seed. In addition, the left anterior-IPL (or l-aIPL) was connected only to the cuneus more in the incongruent condition. Lastly, the right TPJ is connected more in incongruent condition to bilateral STS, temporal pole, and right caudate. See Fig. 5 and Supplementary Fig. 3 for details.

**Figure 5.**
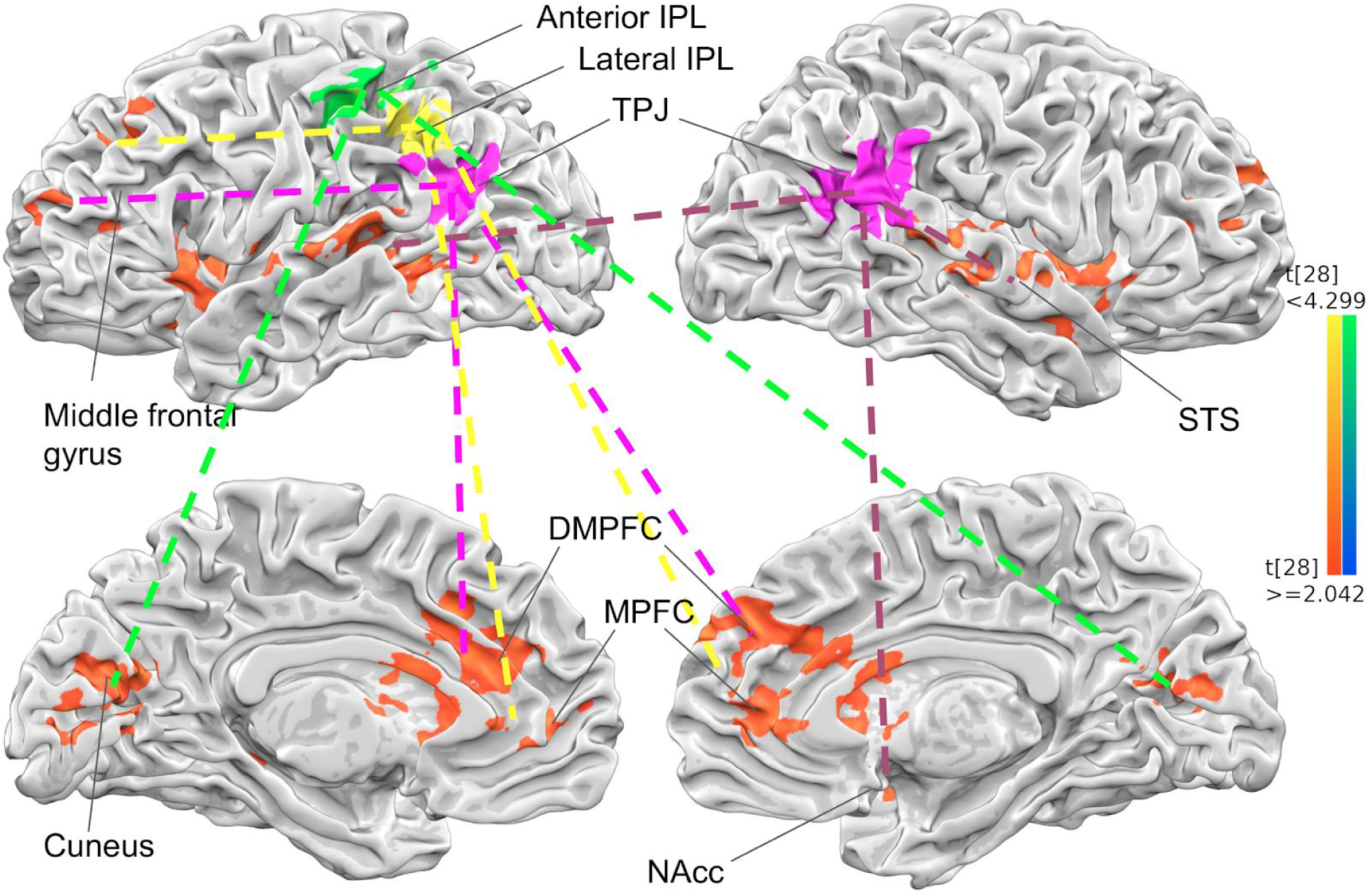
PPI results. The functional connectivity of four different seed regions in ‘incongruent > congruent’ contrast. The left TPJ (pink) are connected to bilateral vmPFC, dmPFC, middle frontal gyrus (pink dotted line), the right TPJ (pink) are connected to bilateral STG/STS and right caudate (violet dotted line), the left lateral IPL (yellow) are connected to bilateral mPFC, dmPFC, left superior frontal gyrus, insular. The anterior IPL (green) was connected to bilateral cuneus. All the surface brain map was shown in p < 0.05, FWE corrected, the PPI was conducted separately for each seed region and concatenated later for visualization, the individual results are provided in supplementary information.

Two outstanding points worth mentioning here: first, maybe partly due to their anatomical adjacency, some of the functional connected networks associated with l-TPJ and l-lIPL are identical, even though both regions are revealed by different statistical analyses (l-TPJ by “congruent vs. incongruent” GLM contrasts; whereas l-lIPL by incongruence-driven behavioral parameters, and higher activity for more yielding to the significant other’s opinion), suggesting that it may be that these two (or even three) neighboring brain regions may share functions related to self-other evaluation and conflict resolving. Second, the bilateral TPJ, which are involved in both “shop together vs. shop alone” and “congruent vs. incongruent” contrasts, showed separate functionally connected networks, a claim which has been been proposed about the separated roles in r-TPJ and in l-TPJ^44^.

Since the two GLM contrasts and two parametric modulations have jointly shown the by-far most outstanding find: the left temporal-parietal brain regions: l-TPJ, l-l IPL, and l-aIPL (Figure 6 left, with colored pink, yellow, and green in the bottom-up order), their inter-connectivity, other than the obvious anatomical adjacency, deserve closer examination. To do so, we implement the mediation analysis among these three ROIs, to test a causal model of whether the effect of l-aIPL activity (the x variable) on subsequent l-TPJ activity (the y variable) is mediated via changes in l-lIPL (the mediator variable)^45^. While the total effect under contrast of “together vs. alone” was significant from l-aIPL to l-TPJ (total *c* = 0.559, *se* = 0.117, *p* < 0.001), such total effect is better explained by the mediation of the l-lIPL (indirect path *a*b* = 0.279, *se* = 0.084, *p* = 0.002), than the direct effect of l-aIPL to l-TPJ (relatively less significant: direct path *c’* = 0.279, *se* = 0.120, *p* = 0.028) (See Fig. 6 for more details). In contrast, the reverse mediation of l-lIPL was not significant in an bottom-up-style model, namely predicting l-aIPL (the y variable) activity from l-TPJ activity (the x variable) (*a*b* = 0.207, *se* = 0.197, *p* = 0.301), although the connectivity between l-TPJ and l-lIPL was still significant (*a* = 0.946, *se* = 0.155, *p* < 0.001) (See the dotted-lines and descriptions in the inner triangle of Fig. 6). Put together, the results reflect the reciprocal interactions among three ROIs during “shop together vs. shop alone” context, by two two-way interactions between l-TPJ and l-lIPL, and between l-TPJ and l-aIPL, but only one-way top-down influence from l-aIPL to l-lIPL. With the presumed underlying functions of l-TPJ (mentalizing significant other), l-lIPL (insisting-to-yielding), and l-aIPL (evaluating self/other-preference), these interactions (shown in Fig. 6) help underpin the complex and dynamic social behaviors with the similarly complicated brain network dynamics.

**Figure 6.**
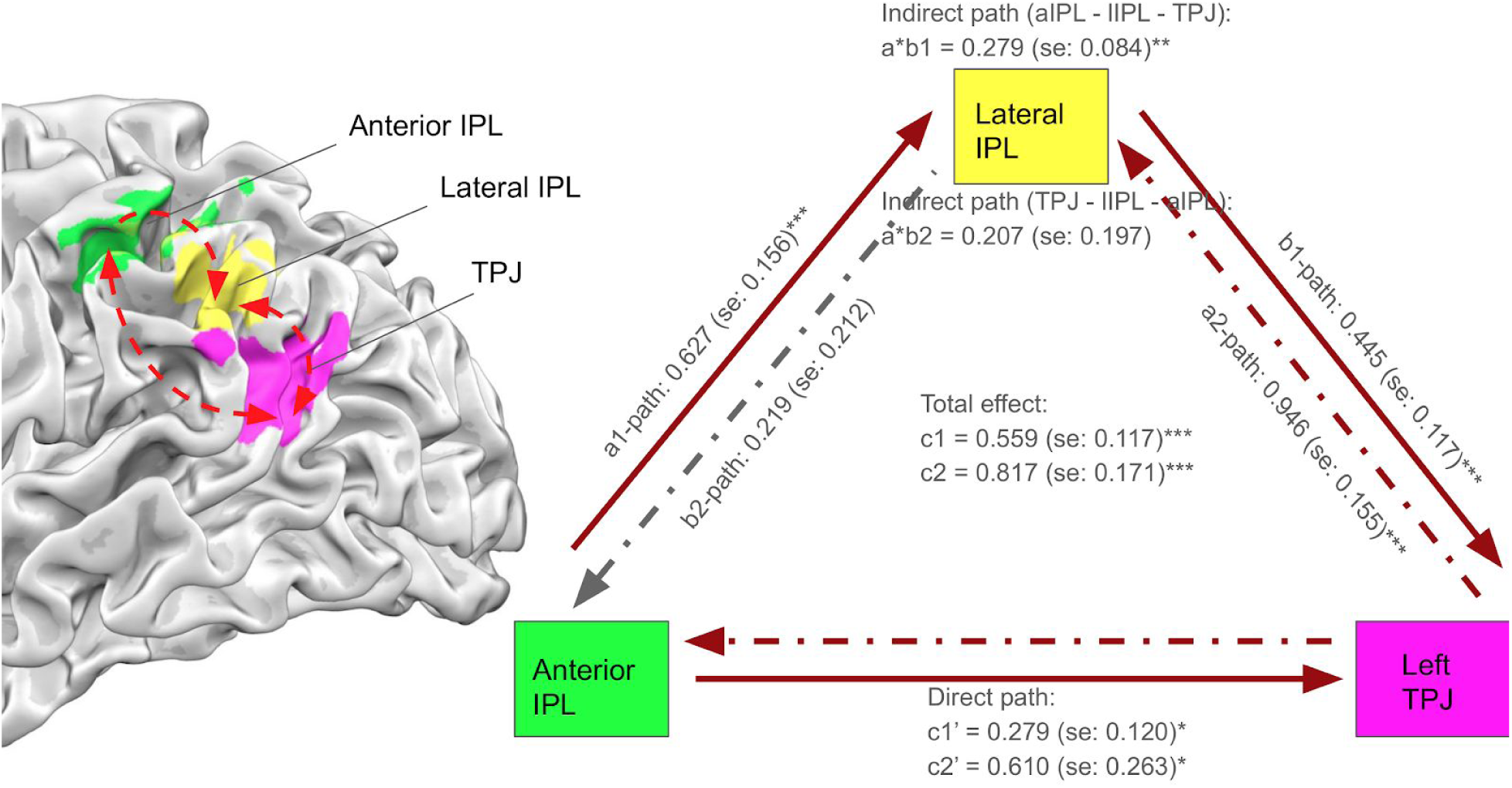
Mediation analysis results. The subject-level effective relationship during ‘together > alone’ between three ROIs, the left anterior IPL (l-aIPL, green), lateral IPL (l-lIPL, yellow), and TPJ (pink) (left), are shown in two different subsequent ways (right). In the causal model from l-aIPL to TPJ (*c1*, outward triangle denoted by lines), or from TPJ to l-aIPL (*c2*, inward triangle denoted by dotted lines), the total effect is significant (*c1* and *c2*, \*\*\**p* < 0.001) as well as the direct path of models (*c1*’ and *c2’*, \**p* < 0.028). However, the mediatory role of l-lIPL (*a*b*) was significant in only one way: from l-aIPL to TPJ (indirect path, *a*b1 (a-lIPL to TPJ)*, \*\**p* = 0.002). Meanwhile, neither effects of mediation (indirect path, *a*b2 (TPJ to a-lIPL), p* = 0.301) nor path from l-lIPL to l-aIPL (*b2, p* = 0.309) was significant in the second causal model (TPJ - l-lIPL - l-aIPL).

## Discussions

This study investigates the neural correlates of preference and decision making of college couples in shopping, and how their decisions are modulated by each other’s item preferences. Previous studies seeking to understand the influence of others in decision making have adopted design/manipulations like: the number of (virtual) responders in consensus decision making^46^, using speech for two-people communications^47^, or even hyperscanning between two scanners simultaneously^48^. What distinguishes our current study from the rest of “above-mentioned” fMRI studies on social interactions is that, first, while prior studies used either 1 scanner at a time, and two separate scans with subjects doing connected tasks (e.g., speech communication), or two scanners with people doing tasks simultaneously, ours is a hybrid of having two people doing together, but with one scanner only. Such arrangements bypass the hurdle of technical challenges of hyperscanning. Second, the introduction of real college couples not only increase the face validity (and therefore appeal) of the current study, but also avoid concern when people are told to respond along with other confederates in the scan. In other words, our study features a real life-like task design in which the participants have the highest guarantee that they are interacting with partners during the experiment.

Along with previous literature^4,7,49^ that suggest the major influencers of individual decision making are from significant people like family members or partners in most social contexts^50,51^, the behavioral results of this study further echo an earlier one^5^ in that couples, or people in the mating mindset (relative to single mindset), tend to comply with partner’ opinions, even in the face of preference conflicting with one’s own. Another interesting find from the current behavioral data, other than the clear modulation of partner’s opinion in one’s shopping decisions, is that 68% of the decisions in the “shop-together” trials are harmonious or congruent, unbeknown to the participants. There are at least two possible explanations, one being the inherent taste of couples (e.g., “birds of a feather flock together”), and another being the nature of the “together” trials favoring this kind of social conformity between couples, a tendency that is “*subtle, indirect, heuristic-based, and* sometimes *outside of awareness*”^52^. While the current two hypotheses cannot be examined without additional behavioral experiment in which the couples do the task “alone” at all times (as the baseline comparison), the lack of “expected” sex differences in the modulation of together-incongruent trials, or the similar yielding/insisting responses between male and female participants, are consistent with this suggestion of ‘implicit social agreement for harmony’ for all participants. After all, the goal of the current study is to create naturally incongruent conditions in which the two-people fMRI can help map neural substrates of social decision making, not to exhaust the reasons behind this seemingly ‘social conformity’, which awaits future research efforts.

The brain areas that are commonly activated by “together vs. alone” and by “congruent vs. incongruent” contrasts: the bilateral TPJ and precuneus (Fig. 3), in which the former, along with nearby brain areas, such as the STG/STS^53^, are commonly associated with various forms of social processing (TOM), from passively viewing (or “mirroring”) others’ actions^54^, to actively inferring others’ goals^54,55^, beliefs^20^, or moral dilemmas^56^. The precuneus, on the other hand, is typically associated more with self-related and spatial frame-of-reference^57^. But it may be hard to tease apart the ambiguity between “me thinking about others” vs. “me simulating how I would be in his/her shoes” when facing the co-activation of TPJ-precuneus in mentalizing others. Other brain areas in Fig. 3, such as both dmPFC^58,59^ and NAcc^60–62^, are more activated in “shop-together” conditions, with dmPFC related to conflict monitoring and detection^63,64^, and NAcc to predicting behavioral changes to a wide variety of rewarding stimuli^65^. Lastly, the anterior and posterior insular are separately more activated to incongruent and congruent trials, respectively, representing the functional dissociation in insular cortex by different degree of empathy, pain^66,67^, or discomfort from misalignment with group or social values^68^.

Two of the left IPL regions, l-aIPL and l-lIPL, were adjacently modulated by parameters of own-high/other-low to own-low/other-high, and high-difference/insisting to high-difference/yielding, indices (both -3 to 3, with 0 both being congruent trials. Fig. 4). Such juxtaposition of l-aIPL and l-lIPL is rarely seen, be it from the literature or from the analysis viewpoints. While most other studies that have adopted parametric modulation and found IPL activities in tasks like detecting incongruence between participants’ motor actions and resulting perception^69^, moral judgment on magnitude of offender’s liability^70^, or the degree of accuracy in viewing one’s own image (of various ages) morphed^71,72^, to our knowledge there are few, if not none, studies adopting two kinds of parametric and found adjacent brain areas in a single report. While IPL has been suggested as one of the most complicated regions subserving higher-order functions^18,20,73,74^ subserving various processes related to self-evaluation^75^ or attention^76^, our left aIPL-lIPL-TPJ findings provide a more detailed map of this hub region, modulated by at least two kinds of evaluation (self preference and self-other relation) and the subsequent decision.

The functional connectivity (FC) results with 4 chosen ROIs (Fig. 5 and Sup. Fig. 3) yielded both similar or disparate patterns of incongruent-driven connected networks: among them the two proximous regions, l-TPJ and l-lIPL, showed common NAcc, mPFC/dmPFC, l-IFG, and l-insula, etc. These similar FC patterns between l-TPJ and l-lIPL seeds, along with the highly significant effective connectivity (i.e., mediation) analysis (Fig. 6), supports their unitary nature of these two seeds and connected networks underpinning social interactions: including empathic choices in l-IFG and l-MFG^21,77^, saliency and discomfort of social disagreements in l-insula^68^, and effort and reward in/after resolving potential conflicts in mPFC and NAcc^78^, etc. Right-TPJ seed, in contrast, showed connection with bilateral STS/STG, reflect efforts in preference comparisons and theory-of-mind^79^. Lastly, the l-aIPL would reasonable to be connected to cuneus (i.e., more visual evaluations) in the incongruent trials, given its responsiveness to ‘self-high/other-low’ trials. All in all, these interconnected ROIs, along with their connected networks, work together to achieve the desired outcome of social harmony.

The line-up of three left IPL-TPJ ROIs, mainly l-aIPL, l-lIPL, and l-TPJ, as well as the two mediation results (see Fig. 6) also have their own implications. For example, the middle l-lIPL seems as a ‘bridge’ of the two neighboring regions: the upper l-aIPL for comparing self(-high) to (the significant) other(-low), and therefore more self-oriented, and the lower l-TPJ for mentalizing others (therefore other-oriented). The two directional mediation analyses, l-aIPL->l-lIPL->l-TPJ and l-TPJ->l-lIPL->l-aIPL, reveal both the significant and reciprocal connectivity between l-TPJ and l-lIPL as a highly connected ‘unit’ for social resolution, apparently reflecting the social desirability of more (68%) congruent trials between couples. In addition, the fact that l-aIPL is influencing more to the l-TPJ (but not the other way around), also speaks to the other-, rather than self-, concerned state of mind in together trials. Together, both the functional (PPI) and effective (mediation) connectivity analyses jointly delineate the dynamic nature of interactions between couples during shopping.

In recent review articles about TPJ-IPL^20,53,73^, we have noticed a growing trend of combining multi-modal methods, including resting-state fMRI, Diffusion Tensor Imaging (DTI), along with multiple task-based fMRI, and with cytoarchitectonic studies, furthering our understanding about structural subdivisions and the functional coverage in this highly complicated higher order brain regions. Paralleling with these aggregational efforts, the current study, likewise, adopts multiple analysis methods (GLM contrast, parametric modulation, PPI, and mediation analyses), providing results of different angles into these TPJ-IPL complex, along with its interconnected networks. In a similar spirit, our study, although narrow^80^, helps address this ubiquitous social phenomenon (interacting college couples), with two-people fMRI experiments, and illuminate inner interactions in IPL-TPJ, on top of their alluded functions, including (but not limited to): attention, memory, language, with social behaviors. Future studies, with the similar vein, should also follow the path toward more transparency in data, stimuli, code, and even scripts for advanced analyses^81^.

## Methods

### Participants

Thirty healthy adults (16 males; mean age=22.7±2.57 yrs, out of the 22 participating couples), who were recruited via online ads and snowball sampling methods, joined the fMRI experiment taking place at the Mind Research and Imaging (MRI) Center, National Cheng Kung University (NCKU). Eligible couples were to be in a romantic relationship for at least a year, with normal or corrected-to-normal vision, and no known neurological disorders. All participants provided written informed consent prior to the study. All methods and procedures were approved by NCKU Governance Framework for Human Research Ethics (number 104-206).

### Stimuli and Procedure

**Stimuli** 463 items and their prices (New Taiwanese Dollars, or NTD) were collected from local online shopping websites (such as ruten, yahoo, pchome, etc). The mean price was 1529±3328 NTD ($1 USD = $30 NTD), ranging from 15 to 45000 NTD. The stimuli were available online at GoogleDrive (https://drive.google.com/drive/folders/1lP76r1zlZfuJuNc2d1xTEKkr6Q9MRKWi?usp=sharing).

### Task

In this event-related fMRI study, two types of shopping contexts were designed to examine how significant others’ preferences affect one’s own shopping decisions. As shown in Fig. 1 (upper left), shopping ‘alone’ trials were indicated by the two inward-pointing, whereas shopping ‘together’ trials outward-pointing, arrows alongside the fixation cross at the beginning of each trial, and both conditions were with 100% cue validity. After the fixation cross and arrows (1s), one stimulus was presented with the price underneath on the screen of the two connected-PCs, with evaluation time up to 6s or until one’s own response, whichever came first. Participants were required to give preference ratings from 1 (dislike most) to 4 (like most) during these 6s period. Then the two screens would show either one own’s rating (always on the left, alone trials), or own-plus-other rankings (on both sides, together trials), together with the stimulus, for a jittered interval (3-6s, or the so-called ‘modulation period’), during which the two evaluations will be transmitted, and presented (for together trials) or kept identical (for alone trials, no modulation here). Subsequently, both a ‘checked’ and a ‘X’ mark would appear in the lowest part of the screen, cueing the beginning for response period (up to 6s). Each participant’s responses would be recorded and displayed independently by a ‘purchased’ or ‘not-purchased’ sign, and would last until the 6s period was up (and followed by fixation cross for variable inter-trial intervals, or ITIs). The trial condition orders and ITIs were determined by optseq2 (https://surfer.nmr.mgh.harvard.edu/optseq/).

Each participant was scanned for 4 or 5 runs, and each ∼5 min run contained 20 to 25 trials (trial length: 16-19s). For the first 5 subjects, the trial distributions between alone and together trials were half-half. But to increase the number of together trials, for the remaining 25 participants, the percentage of together trials were raised to 2/3 (and alone trials ⅓).The experiment and internet communications were programmed via Psychtoolbox^82^ and TCPIP plugins for Matlab.

### Procedure

Upon completing informed consent, the couple was told that the experiment was like an online shopping context they experience everyday: imagine you have a credit card with enough credit amount, so that you can buy whatever you like or fits you (but you typically will not buy them all), and the typical trial consisted of: beholding an item, rated how much you like it, then with occasional rating from the significant other alongside, and then make your final decisions. Then one of romantic partners volunteered in the scanner, while the other remained in the waiting room, facing the internet-connected desktop PC to interact with the one inside the scanner. The couples were then familiarized with two kinds of trial format, and practiced until both understanding were assured. Eight (out of 22) pairs were able to do consecutive scans (e.g., boy first, girl second) in the same day. After the fMRI, the couple were reimbursed for 1200∼1400 NTD for their time (usually 1-2 hrs each).

## Data acquisition, preprocessing, and analysis

### fMRI data acquisition and preprocessing

Images were acquired with a 3T General Electric 750 MRI scanner (GE Medical Systems, Waukesha, WI) at the NCKU MRI center, with a standard 8-channel head coil. Whole-brain functional scans were acquired with a T2* EPI (TR= 2 s, TE= 35 ms, flip angle = 76 degree, 40 axial slices, voxel size = 3 × 3× 3 mm^3^). High-resolution structural scans were acquired using a T1-weighted spoiled grass (SPGR) sequence (TR=7.5s, TE=7.7ms, 166 sagittal slices, voxel size = 1× 1× 1 mm^3^). The fMRI data were preprocessed and analyzed using BrainVoyagerQX v. 2.6 (Brain Innovation, Maastricht, The Netherlands) and NeuroElf v1.1 (http://neuroelf.net). After slice timing correction, functional images were corrected for head movements using the six-parameter rigid transformations, aligning all functional volumes to the first volume of the first run. Neither high-pass temporal filtering nor spatial smoothing was applied. The resulting functional data were co-registered to the anatomical scan via initial alignment (IA) and final alignment (FA), and then both fmr and vmr files were transformed into Talairach space^83^.

### Behavioural data analysis

The analysis of variance (ANOVA) was used to show significant differences between buying ratio across two different sessions (alone/together). The two logistic regression models were separately run by including age, sex, price, and item evaluations (own rating for shop-alone condition, and own rating+other rating/rating difference in the shop-together condition), to predict the binary shopping decisions: buy or not-buy.

### fMRI data analysis

General linear model (GLM) and the specific contrasts of interest were applied to identify associated brain regions under various conditions. Together trials were defined by combining the modulation (upon the presentation of both preferences) and subsequent decision time, and the same summation was applied also to the alone trials (‘no modulation’ + ‘decision’ times). The “together vs. alone” condition was intending to show the main effect of social ‘interaction’, therefore the typical theory-of-mind/TOM regions, such as bilateral TPJ, was expected. In addition, together trials could further be subdivided into ‘congruent’ and ‘incongruent’ conditions, with the former defined as ‘item preferences were both high (3, 4) or both low (1,2)’; and the latter as ‘either one was high (3, 4) and the other low (1, 2), or vice versa (1 or 2 vs. 3 or 4)’. This “congruent vs. incongruent” contrast was expected to reveal brain regions involved in rating harmony (reward), less conflict, less TOM, and then the decision to yield or to insist.

In addition to GLM contrast, parametric modulation effects were further defined as the signed differences between rating differences. For example, in the own-high(/other-low) condition, such as ‘I 4 and other 1’, the difference was +3; or in the other-high(/own-low) condition, such as ‘I 2 and other 4’, the difference was -2. And zero would represent all congruent trials, in the present analysis including ratings of ‘11’, ‘22’, ‘33’, ‘44’, ‘12’, ‘21’, ‘34’, and ‘43’. As for the second way of parameterization, we define the behavioural changes in shopping decision as yielding: giving in to make significant other happy; and insisting: sticking to one’s own preference and ignore incongruent rating from partner. The degree of change was further parameterized by combining the difference between preferences (1 ∼ 3) and decision making (positive for yielding, negative for insisting), resulting in ‘-3’ as most insisting, such as ‘I was 4 and the other 1, and I decided to buy’, or vice versa, like ‘I was 1 and the other 4, and I decided not to buy. Zero was the congruent trials as exactly defined in the 1st parametric analysis. ‘+3’ was the most yielding, such as ‘I was 4 and the other 1, and I decided not-buy’, or vice versa, like ‘I was 1 and the other 4, and I decided to buy’. These two types of parameters were separately incorporated into the GLM, to identify brain regions linearly correlated with various parameters.

Lastly, the psychophysiological interaction (PPI) analysis^35^ was to examine functional connectivity on a priori region of interest (ROI). Four ROIs: the right TPJ, the left TPJ, the left anterior IPL and lateral IPL, revealed by the GLM contrast of ‘together vs. alone’ trials and parametric modulation, were independently downloaded by the term ‘TPJ’ and ‘IPL’ into the online database Neurosynth (http://neurosynth.org/). These 4 seed-based PPIs were all comparing “congruent vs. incongruent” trial conditions (see Fig. 5). The following mediation analysis was carried out to identify directional relations between anatomically proximated ROIs located in left temporal-parietal brain regions. The anterior IPL defined from earlier parametric modulation of preference differences, and meta-analysis derived IPL and TPJ are entered into path analysis as covariates. The connectivity of three regions in the contrast of together > alone are measured in random effect, and compared to null-hypothesis derived by 10,000 times of bootstrapping (Monte Carlo simulation) method.

All group analyses were conducted in random effects models. For the visualization of statistical maps, all the statistical tests were thresholded at *p* < 0.05 (FWE corrected), using a cluster threshold defined by various uncorrected *p* values (0.05, 0.01, 0.005, or 0.001) with 1000 Monte Carlo simulations onto the estimated smoothing kernel size, with the alphasim command under Neuroelf. The anatomical labels of peak voxel in significant clusters were determined by Talairach atlas (tdclient), also implemented in the NeuroElf toolbox. The reported coordinates are in the standard Montreal Neurological Institute (MNI) space.

## Acknowledgments

This work is supported by MOST 105-2420-H-006-002-MY2. We thank Mind Research and Imaging Center (MRIC) at National Cheng Kung University for consultation and instrument availability.

## Supplementary information

**Supplementary table S1.** Behavioral logistic regression. The logistic regression model estimates buying decision (buy/not buy) in alone and together condition, revealing that own preference, price, and age are factors that significantly predict one’s behaviour in both “shop alone” and “shop together” conditions. Additionally, in “shop together” trials, the differences between the preferences of own and other showed significantly negative correlation, indicating the modulatory role of significant others’ preferences in one’s own shopping decisions.

|  | Estimate | SE | tStat | pValue |  | Estimate | SE | tStat | pValue |
| --- | --- | --- | --- | --- | --- | --- | --- | --- | --- |
| Alone | — | — | — | — | Together | — | — | — | — |
| (Intercept) | -2.4475 | 0.87615 | -2.7934 | 0.0052 | (Intercept) | -3.7678 | 0.66254 | -5.6869 | 0.0000 |
| self_rank | 2.4874 | 0.13897 | 17.898 | 0.0000 | self_rank | 2.801 | 0.12679 | 22.092 | 0.0000 |
| price | -0.00035<br>493 | 6.35E-05 | -5.5912 | 0.0000 | ranking_diff | -0.97043 | 0.18667 | -5.1986 | 0.0000 |
| sex_female | -0.03104<br>5 | 0.17426 | -0.17815 | 0.85861 | price | -8.55E-0<br>5 | 2.24E-05 | -3.8175 | 0.0001 |
| age | -0.18324 | 0.036711 | -4.9913 | 0.0000 | sex_female | -0.13655 | 0.13176 | -1.0364 | 0.30001 |
|  |  |  |  |  | age | -0.16101 | 0.027318 | -5.894 | 0.0000 |
|  |  |  |  |  | self_rank:diff | 0.01866 | 0.065315 | 0.28569 | 0.77511 |

**Supplementary table S2.** GLM contrast cluster table. The GLM contrast cluster table in the two contrasts: “Together > Alone” and “Congruent > Incongruent”. All brain coordinates are reported in the standard MNI space, and the contrasts are thresholded at p < 0.01 (Top) and p < 0.05 (Bottom). The cluster threshold was determined by AlphaSim in Neuroelf, using smoothness parameters estimated from the residuals of the statistical map.

| Cluster table of map: "Together > Alone" |  |  |  |  |
| --- | --- | --- | --- | --- |
| x, y, z | k | max | mean | tdclient |
| ----- |  |  |  |  |
| -12, 56, 27 | 959 | 7.040407 | 3.553929 | LH Superior Frontal Gyrus (Brodmann area 9) (-12; 58; 27) [d=2.0mm] |
| -43, -58, -8 | 1169 | 6.532799 | 3.44449 | LH Fusiform Gyrus (Brodmann area 37) (-41; -60; -11) [d=4.1mm] |
| 33, 16, -11 | 109 | 6.304917 | 3.499791 | RH Inferior Frontal Gyrus (Brodmann area 47) |
| 12, -85, 6 | 463 | 6.013853 | 3.416915 | RH Cuneus (Brodmann area 17) |
| -3, -50, 25 | 205 | 5.886473 | 3.493299 | LH Posterior Cingulate (Brodmann area 23) |
| -39, 17, -10 | 309 | 5.851212 | 3.425125 | LH Inferior Frontal Gyrus (Brodmann area 47) |
| -49, 7, 41 | 88 | 5.274084 | 3.258311 | LH Middle Frontal Gyrus (Brodmann area 8) (-50; 7; 41) [d=1.0mm] |
| 53, -61, 22 | 283 | 5.165014 | 3.290112 | RH Middle Temporal Gyrus (Brodmann area 39) |
| 55, -1, -15 | 88 | 5.131149 | 3.368571 | RH Middle Temporal Gyrus (Brodmann area 21) ( 57; -3; -15) [d=2.8mm] |
| 19, 5, 11 | 97 | 4.97557 | 3.379958 | RH Lentiform Nucleus (Putamen) ( 20; 4; 11) [d=1.4mm] |
| 30, -80, 8 | 71 | 4.644316 | 3.138276 | RH Middle Occipital Gyrus (Brodmann area 19) ( 32; -83; 8) [d=3.6mm] |
| 50, 2, 39 | 76 | 4.283547 | 3.184655 | RH Middle Frontal Gyrus (Brodmann area 6) |
| Cluster table of map: "Congruent > Incongruent" |  |  |  |  |
| x, y, z | k | max | mean | tdclient |
| ----- |  |  |  |  |
| 9, -78, 27 | 1309 | 4.855926 | 2.582847 | RH Cuneus (Brodmann area 18) |
| 28, 30, 12 | 200 | 4.706691 | 2.641261 | RH Insula (Brodmann area 13) ( 31; 22; 12) [d=8.5mm] |
| -31, 15, 31 | 167 | 4.503531 | 2.611587 | LH Middle Frontal Gyrus (Brodmann area 9) (-35; 15; 31) [d=4.0mm] |
| -61, -30, 15 | 184 | 4.14919 | 2.558136 | LH Superior Temporal Gyrus (Brodmann area 42) (-61; -28; 15) [d=2.0mm] |
| 37, -20, 8 | 142 | 3.892134 | 2.526385 | RH Insula (Brodmann area 13) ( 38; -20; 8) [d=1.0mm] |
| 39, 7, 29 | 213 | -5.91709 | -2.715866 | RH Inferior Frontal Gyrus (Brodmann area 9) ( 39; 5; 29) [d=2.0mm] |
| 0, 13, 42 | 212 | -4.760147 | -2.554845 | LH Medial Frontal Gyrus (Brodmann area 6) ( -1; 13; 43) [d=1.4mm] |
| -45, 13, 26 | 117 | -4.220338 | -2.646572 | LH Middle Frontal Gyrus (Brodmann area 9) |
| 31, -68, -6 | 113 | -3.504229 | -2.371885 | RH Fusiform Gyrus (Brodmann area 19) ( 31; -68; -7)<br>[d=1.0mm] |

**Supplementary figure S3.**
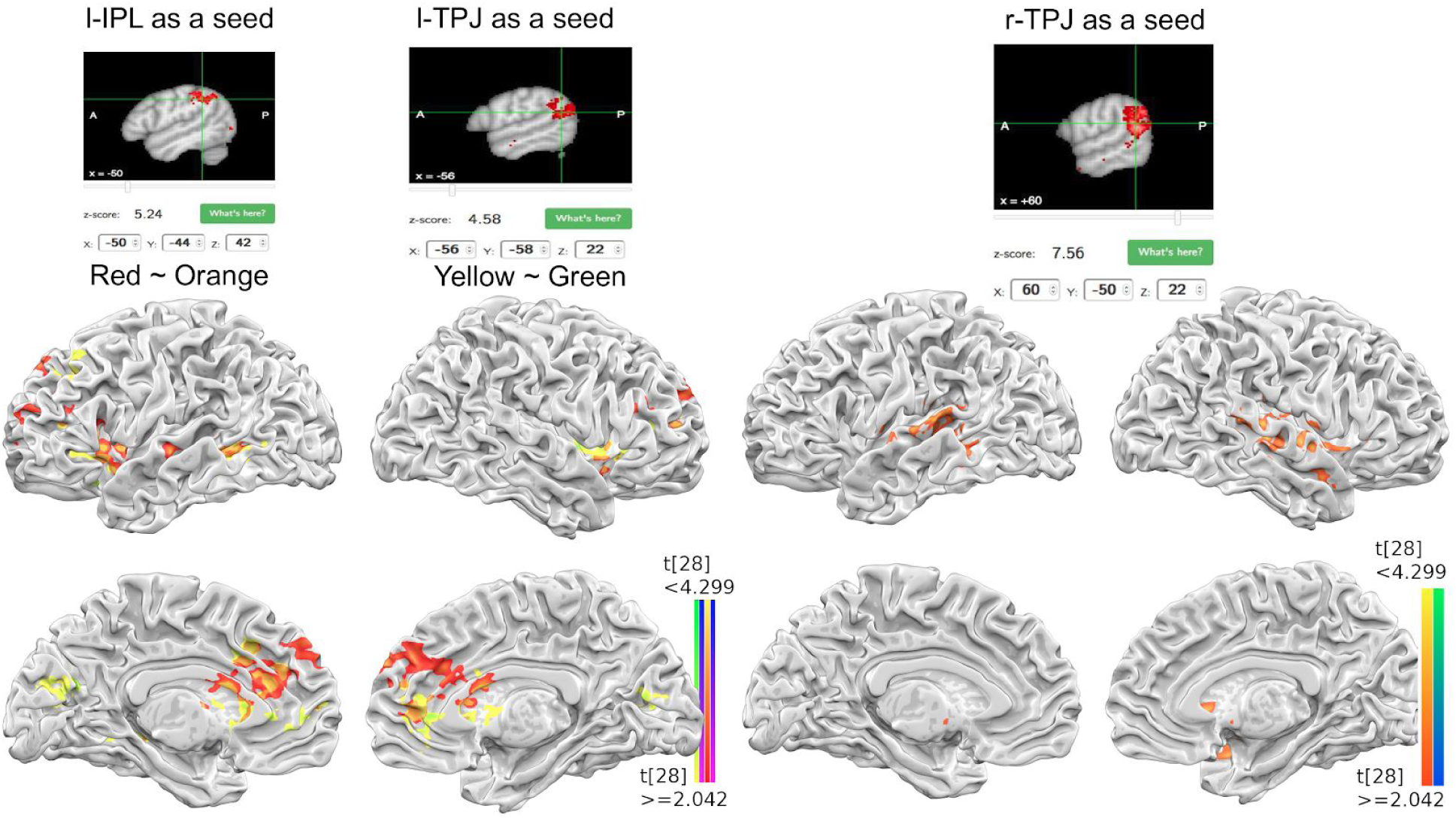
PPI results of three different seed regions shown in separate. The similar functional connectivity between l-IPL and l-TPJ seed were overlaid (left), while the distinct functional connectivity map was shown with r-TPJ seed (right). All brain maps are thresholded at p < 0.05, FWE corrected.

